# Directed Evolution of Aptamer Discovery Technologies

**DOI:** 10.1101/2021.11.23.469732

**Authors:** Diana Wu, Chelsea K.L. Gordon, John H. Shin, Michael Eisenstein, H. Tom Soh

## Abstract

**Conspectus:** Although antibodies are a powerful tool for molecular biology and clinical diagnostics, there are many emerging applications for which nucleic acid-based aptamers can be advantageous. However, generating high-quality aptamers with sufficient affinity and specificity for biomedical applications is a challenging feat for most research laboratories. In this *Account*, we describe four techniques developed in our lab to accelerate the discovery of high quality aptamer reagents that can achieve robust binding even for challenging molecular targets. The first method is particle display, in which we convert solution-phase aptamers into aptamer particles that can be screened via fluorescence-activated cell sorting (FACS) to quantitatively isolate individual aptamer particles based on their affinity. This enables the efficient isolation of high-affinity aptamers in fewer selection rounds than conventional methods, thereby minimizing selection biases and reducing the emergence of artifacts in the final aptamer pool. We subsequently developed the multi-parametric particle display (MPPD) method, which employs two-color FACS to isolate aptamer particles based on both affinity and specificity, yielding aptamers that exhibit excellent target binding even in complex matrices like serum. The third method is a click chemistry-based particle display (click-PD) that enables the generation and high-throughput screening of “non-nattural” aptamers with a wide range of base modifications. We have shown that these base-modified aptamers can achieve robust affinity and specificity for targets that have proven challenging or inaccessible with natural nucleotide-based aptamer libraries. Lastly, we describe the non-natural aptamer array (N2A2) platform, in which a modified benchtop sequencing instrument is used to characterize base-modified aptamers in a massively parallel fashion, enabling the efficient identification of molecules with excellent affinity and specificity for their targets. This system first generates aptamer clusters on the flow-cell surface that incorporate alkyne-modified nucleobases, and then performs a click reaction to couple those nucleobases to an azide-modified chemical moiety. This yields a sequence-defined array of tens of millions of base-modified sequences, which can then be characterized in a high-throughput fashion. Collectively, we believe that these advancements are helping to make aptamer technology more accessible, efficient, and robust, thereby enabling the use of these affinity reagents for a wider range of molecular recognition and detection-based applications.

## Introduction

Antibodies are one of the foundational tools of molecular biology and biomedical research, but nucleic acid-based aptamers offer some critical advantages relative to protein-based affinity reagents. Aptamers are readily chemically synthesized in a reproducible manner and can be engineered to incorporate useful functionalities such as structure switching^1^ and selective response to environmental conditions such as pH^2^ or redox states^3^. Furthermore, aptamers can be generated for low-molecular-weight targets that would be challenging for antibody generation, such as small-molecule ligands^4,5^ and even metal ions^6^. These attributes make aptamers particularly promising for applications such as biosensors or controlled drug delivery. Aptamers are selected based on their target recognition via an *in vitro* process known as systematic evolution of ligands by exponential enrichment (SELEX).^7,8^ This procedure has three main steps: 1) binding of single-stranded nucleic acids to the target, 2) partitioning of bound nucleic acids from unbound nucleic acids, and 3) PCR amplification of nucleic acids with affinity for the target as a prelude to either further screening or functional characterization. After SELEX, the DNA pools are subjected to high-throughput sequencing (HTS) to identify highly enriched sequences that are likely to have undergone selection due to their target binding. Finally, promising sequences identified from these data are characterized in assays that enable quantification of their target affinity and specificity.

However, the SELEX process is also prone to many challenges. At a high level, this is because the success or the failure of an aptamer discovery campaign can only be assessed at the end of the selection process, and the researcher cannot quantitatively monitor the quality of the aptamer pool (in terms of affinity and specificity to the target) - during the selection process. Over the past decade, our group has made a systematic effort to overcome this crippling limitation and we have developed four technologies (**Figure 1**) that we believe represent particularly important and enabling tools for the successful generation of useful, high quality aptamers.

**Figure 1.**
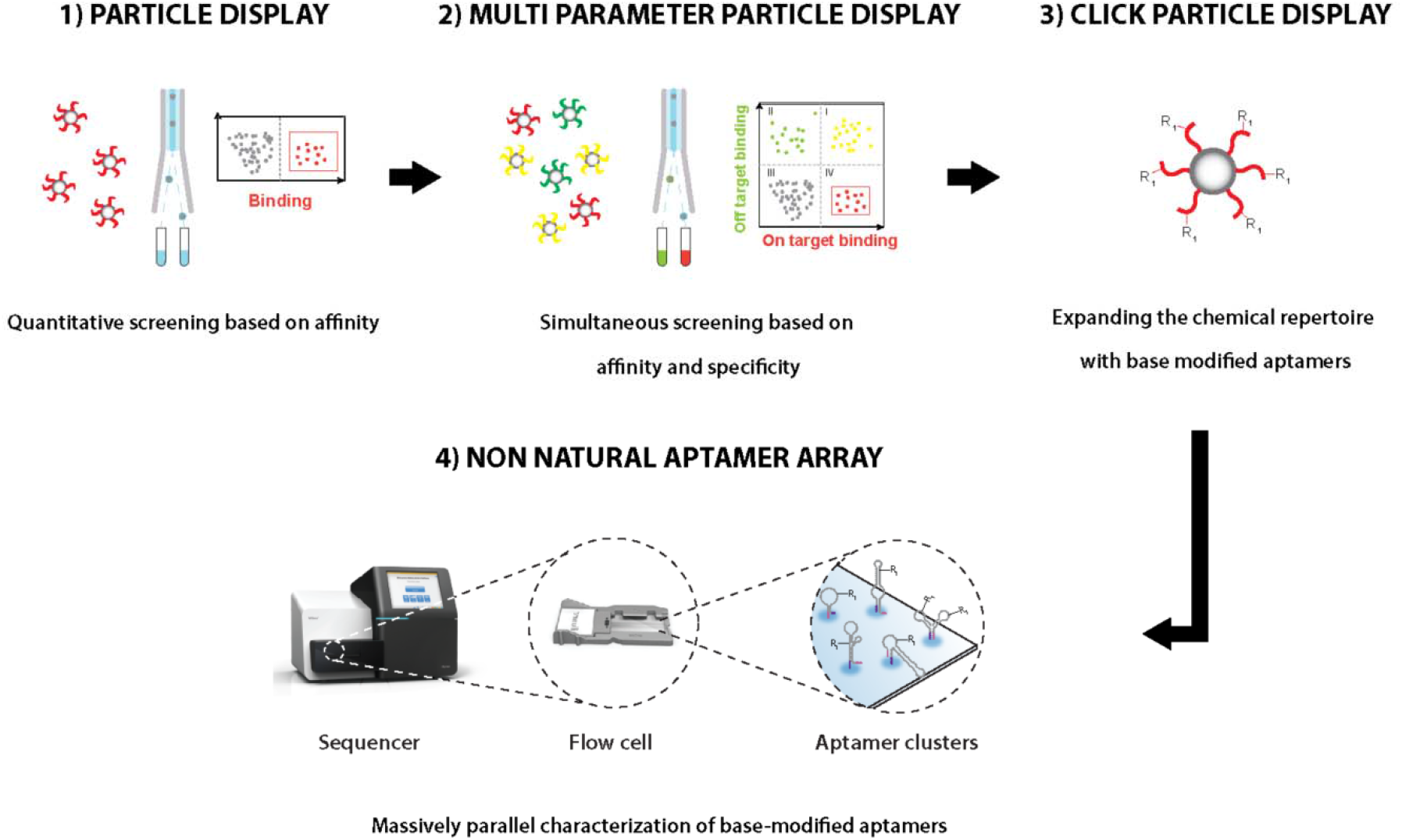
Four aptamer generation and characterization technologies developed by the Soh Lab. 1) Particle display, which allows us to enrich aptamers faster. 2) Multi-parameter particle display, which allows us to generate aptamers that simultaneously achieve high specificity and affinity. 3) Click particle display, which allows us to generate aptamers with expanded chemical diversity. 4) Non-natural aptamer array, which allows us to characterize millions of base-modified aptamers simultaneously.

One of the greatest limitations on conventional SELEX is the number of rounds required to converge the vast starting library of random sequences to a pool of high-quality aptamer candidates. This can require as many as 8–20 rounds of SELEX^9,10^ due to the inherent limitations on the efficiency of aptamer enrichment that can be achieved in a single round, creating ample opportunities for the introduction of PCR biases and other artifacts that can degrade the overall quality of the selected pool. To address this, we developed our particle display platform,^11^ which makes it possible to individually screen large numbers of aptamer candidates in a high-throughput manner using fluorescence-activated cell sorting (FACS). In this approach, aptamer libraries are converted to monoclonal ‘aptamer particles’, which are then screened on the basis of their binding to a fluorescently-labeled target, enabling the rapid selection of high-affinity aptamers in as few as three or four rounds—even for challenging targets that have proven recalcitrant against screening in prior efforts.

We have subsequently identified several strategies for further extending the core particle display technology to enable the discovery of aptamers with improved binding properties. First, in order to isolate aptamers with excellent specificity for their target, we developed multi-parameter particle display (MPPD). Here, we incubate our aptamer particles with differentially fluorescently-labeled target and non-target molecules, and then use two-color FACS to rapidly discriminate aptamers that exclusively bind the target with high affinity.^12^ This approach yields aptamers that achieve far better specificity than those isolated via conventional SELEX in buffer, and we have demonstrated that MPPD-derived aptamers can maintain robust target affinity in complex biological samples such as serum. We next developed click particle display (click-PD)^13^—a strategy to expand the chemical repertoire of natural DNA and RNA, in which we combined particle display with a click chemistry approach to generate aptamer libraries containing nucleobases that can be modified with virtually any chemical moiety of interest. This is valuable, as there is evidence that the limited chemical diversity of natural DNA relative to proteins restricts the range of targets that are suitable for aptamer generation, particularly in the context of hydrophobic proteins and small molecules.^14^

Most recently, in a departure from our particle display-based efforts, we have developed the non-natural aptamer array (N2A2) platform. This is a semi-automated system based on a modified benchtop HTS machine that integrates high-throughput generation, sequencing, and characterization of base-modified aptamers in a single instrument. N2A2 makes it easier to perform aptamer characterization, which has remained one of the most time-consuming and low-throughput aspects of the aptamer-generation process. Importantly, N2A2 enables the rapid characterization of both natural DNA and click chemistry-modified non-natural aptamer sequences with excellent affinity and specificity. In the following sections, we describe in detail these various technologies and the capabilities that they offer in terms of accelerating and improving the efficiency of the aptamer screening process—thereby extending the utility and value of aptamers as a research and clinical tool.

### Accelerating aptamer discovery with particle display

One of the major constraints on conventional SELEX is the efficiency with which high-affinity candidates can be partitioned from low-affinity or non-binding sequences, necessitating many rounds of screening for each aptamer selection. Particle display was developed as a solution to this problem^11^ (**Figure 2**). Particle display begins with a library of aptamer candidates that has been pre-enriched for affinity to the target through a few rounds of conventional SELEX to reduce the number of sequences to a scale that can realistically be analyzed in a single FACS experiment. This pre-enriched library is then converted to monoclonal aptamer particles that each display ~10^5^ copies of the same sequence via an emulsion PCR process, where the reaction can be tuned to ensure that each droplet contains no more than one template. The droplets are subjected to PCR amplification, after which the aptamer particles are recovered and incubated with fluorescently-labeled target molecules, such that the affinity of each aptamer sequence can be directly measured using FACS. The fluorescence intensity of each aptamer particle is directly proportional to the target affinity of the displayed aptamer, and the particles with the highest fluorescence can thus be isolated and PCR amplified to produce the starting pool for the next round. Once there is no longer meaningful enrichment of the target-binding subpopulation of aptamer particles, the pool is sequenced to identify aptamer candidates for further characterization.

**Figure 2.**
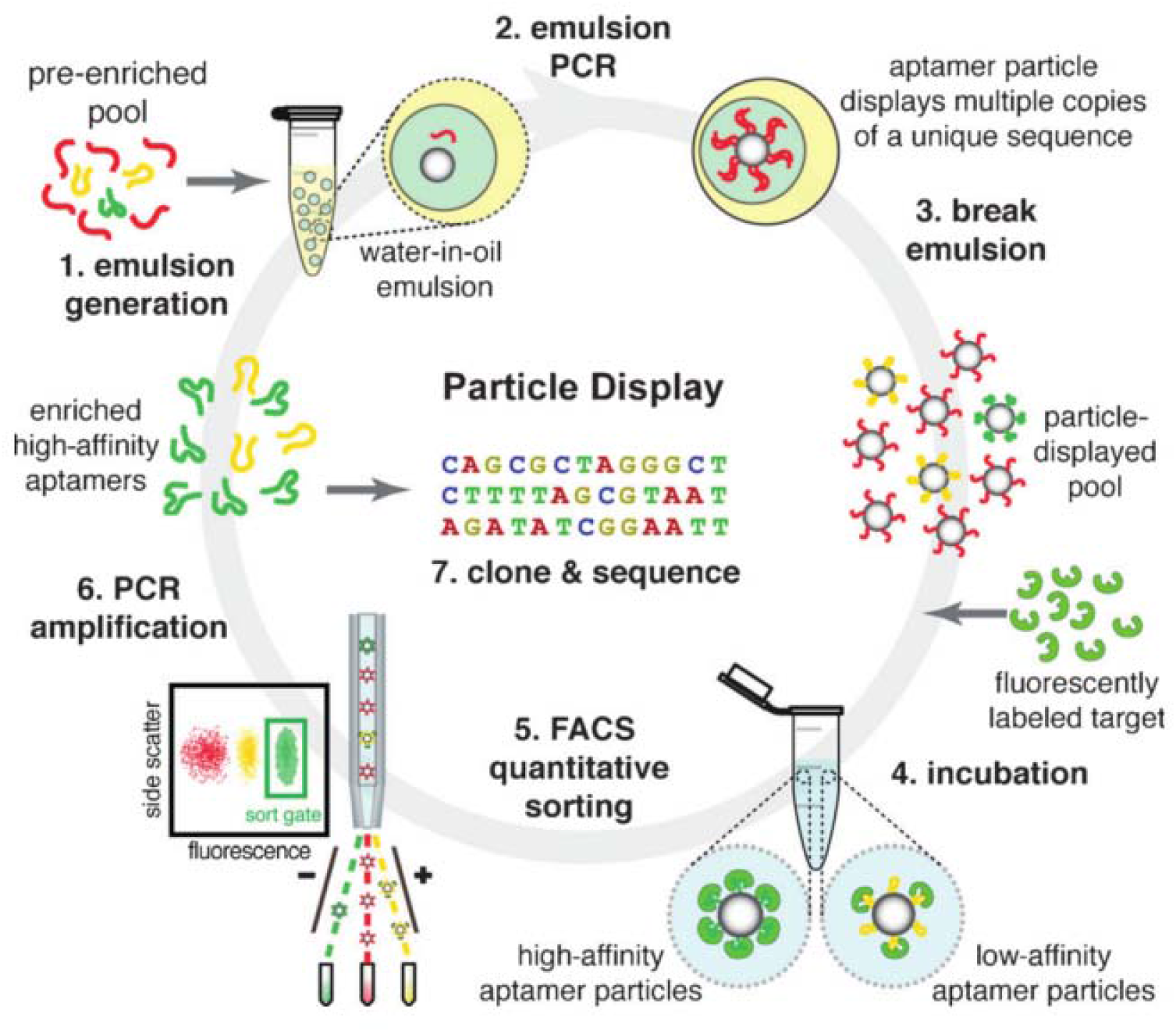
Overview of particle display. 1) A pre-enriched aptamer pool is converted to monoclonal aptamer particles using 2) emulsion PCR. The aptamer particles are 3) isolated and then 4) incubated with fluorescently-labeled target, after which aptamer particles with strong fluorescent signal are 5) collected using fluorescence-activated cell sorting (FACS). The selected sequences are then 6) PCR amplified as a prelude to another round of screening, or 7) sequencing to characterize the resulting pool once the level of enrichment reaches a plateau.

This process results in efficient aptamer partitioning. Mathematical modeling of the theoretical maximum enrichment that can be achieved with conventional SELEX has indicated that any given target-binding sequence will only be enriched by 100- to 1,000-fold in a single round^15,16^. Accordingly, a typical SELEX experiment requires 8–20 rounds to converge the aptamer pool towards a set of sequences with high affinity for the target. However, this also increases the risk of lowering the pool quality through emergence of biases and artifacts, such as incorrectly-sized byproducts^17^ or lower library diversity^18^. In contrast, particle display enables high-throughput screening of the binding properties of large numbers of individual aptamer sequences in each round rather than simply subjecting the pool to a bulk selection process, making it much more efficient to identify and isolate high-affinity binders. We have calculated that an aptamer with an equilibrium dissociation constant (*K*_*D*_) of 100 pM within a pool of aptamers with an overall *K*_*D*_ of 1 nM can theoretically be enriched 1.7 × 10^9^-fold in a single round of particle display screening.^11^ This enables isolation of higher affinity aptamers in far fewer rounds of screening, and also prevents unwanted enrichment of low-affinity sequences based on stochastic binding events, which can occur in conventional SELEX screens. These rare events do not produce a strong enough fluorescent signal for FACS separation to occur, and only the strong signal generated by the binding of multiple labeled target molecules to multiple aptamer copies on a given particle will register as true positives. FACS also makes it possible to directly visualize the progress of the screen and adjust the stringency of screening from round to round, giving users a level of control and insight into the screening process that is not available with standard SELEX.

We have used particle display to rapidly isolate high-quality DNA aptamers for a diverse range of proteins.^11^ As described above, the use of FACS made it possible for us to visually observe the target-binding fraction of the aptamer pool in each round (**Figure 3A**) and adjust the sort gates to isolate only those particles that clearly exhibit high fluorescence (*i.e.,* strong target binding). We also decreased the target concentration from round to round, resulting in more stringent selection of the highest-affinity aptamers. Using this process, we were able to isolate high-quality DNA aptamers for four different targets in just three rounds. We identified aptamers for thrombin and ApoE that exhibited far superior affinity to previously generated aptamers.^19–21^ For example, our thrombin aptamer exhibited a *K*_*D*_ of ~7 pM (**Figure 3B**), which is three orders of magnitude lower than that of previously published thrombin aptamers (*K*_*D*_ = 2.6–5.4 nM). We also generated the first reported natural DNA aptamers for a pair of challenging protein targets, PAI-1 and 4-1BB, with *K*_*D*_s of 339 pM and 2.32 nM, respectively—comparable to previously-reported aptamers that incorporated non-natural bases (*K*_*D*_ = 200 pM and 4 nM, respectively).^22^

**Figure 3.**
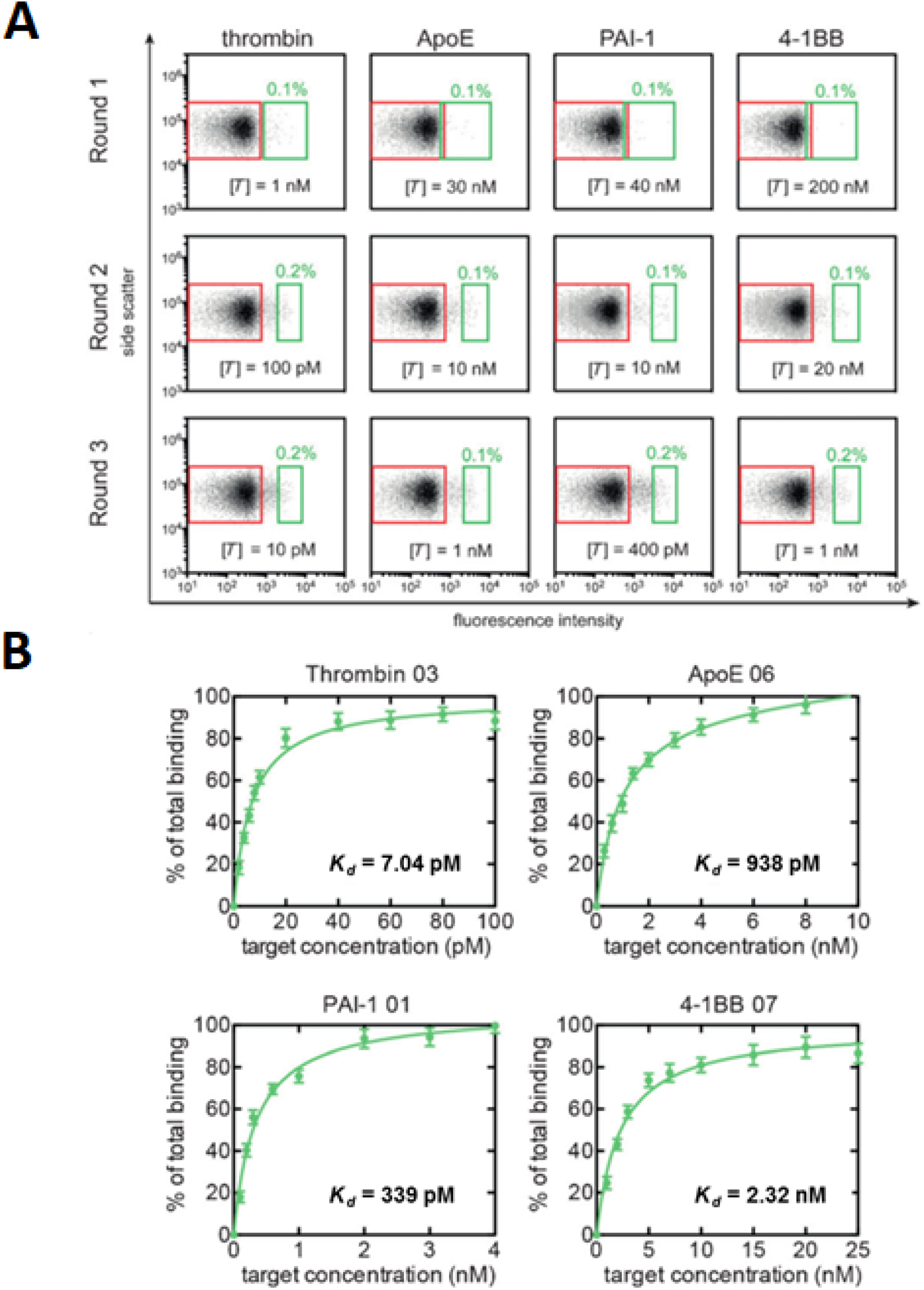
Particle display screening for four different protein targets. **A**) Reference (red) and sort (green) gates for multiple rounds of FACS analysis of particle display screens for thrombin, ApoE, PAI-1, and 4-1BB. In each round, a larger proportion of the aptamer particles binds to the target despite decreasing the target concentration. **B**) Binding curves for the highest affinity aptamers from our particle display screening experiments.

### Selecting for specificity with multi-parametric particle display (MPPD)

Although affinity is a critical parameter for aptamers, it is also essential that the isolated sequences exhibit robust specificity and minimal off-target binding. This is particularly important for the detection of analytes in complex samples such as serum, where promiscuous binding can produce false results in an assay. One widely used solution to this problem entails following standard SELEX with additional rounds of a ‘counter-SELEX’ procedure^23^ that eliminates aptamers with cross-reactivity to non-target or interferent molecules. This approach can be effective, but also creates opportunities to unwittingly discard aptamers that strike an optimal balance between affinity and specificity. This is because each cycle of SELEX is focused entirely on optimizing only a single aptamer parameter at a time – either affinity or specificity – rather than maintaining continuous selection pressure for both characteristics throughout the screening process. Furthermore, counter-SELEX requires even more rounds of PCR amplification and screening, which further exacerbates the risk of PCR biases and artifacts.

To address this problem, we developed MPPD—a particle display-based strategy that allows us to select for both characteristics simultaneously by exploiting the multi-color sorting capabilities of the FACS instrument.^24^ In each MPPD experiment, target and non-target proteins are each labeled with fluorophores that emit at distinct wavelengths. During FACS analysis, we can then readily detect and isolate only those aptamers that exhibit both strong target-associated fluorescence (high affinity) and low non-target-associated fluorescence (high specificity) (**Figure 4A**). We used MPPD to screen for DNA aptamers for tumor necrosis factor α (TNF-α) in a background of diluted human serum, where TNF-α was labeled with AlexaFluor-488 (green) and the serum proteins were non-specifically labeled with AlexaFluor-647 (red). By screening for aptamers with high green and low red fluorescence (**Figure 4A**, quadrant IV), we could enrich aptamers with high TNF-α affinity and minimal affinity for background proteins in just four rounds. In order to demonstrate the added specificity conferred by MPPD screening, we also performed a conventional particle display screen for TNF-α in buffer, where selection was exclusively based on affinity.

**Figure 4.**
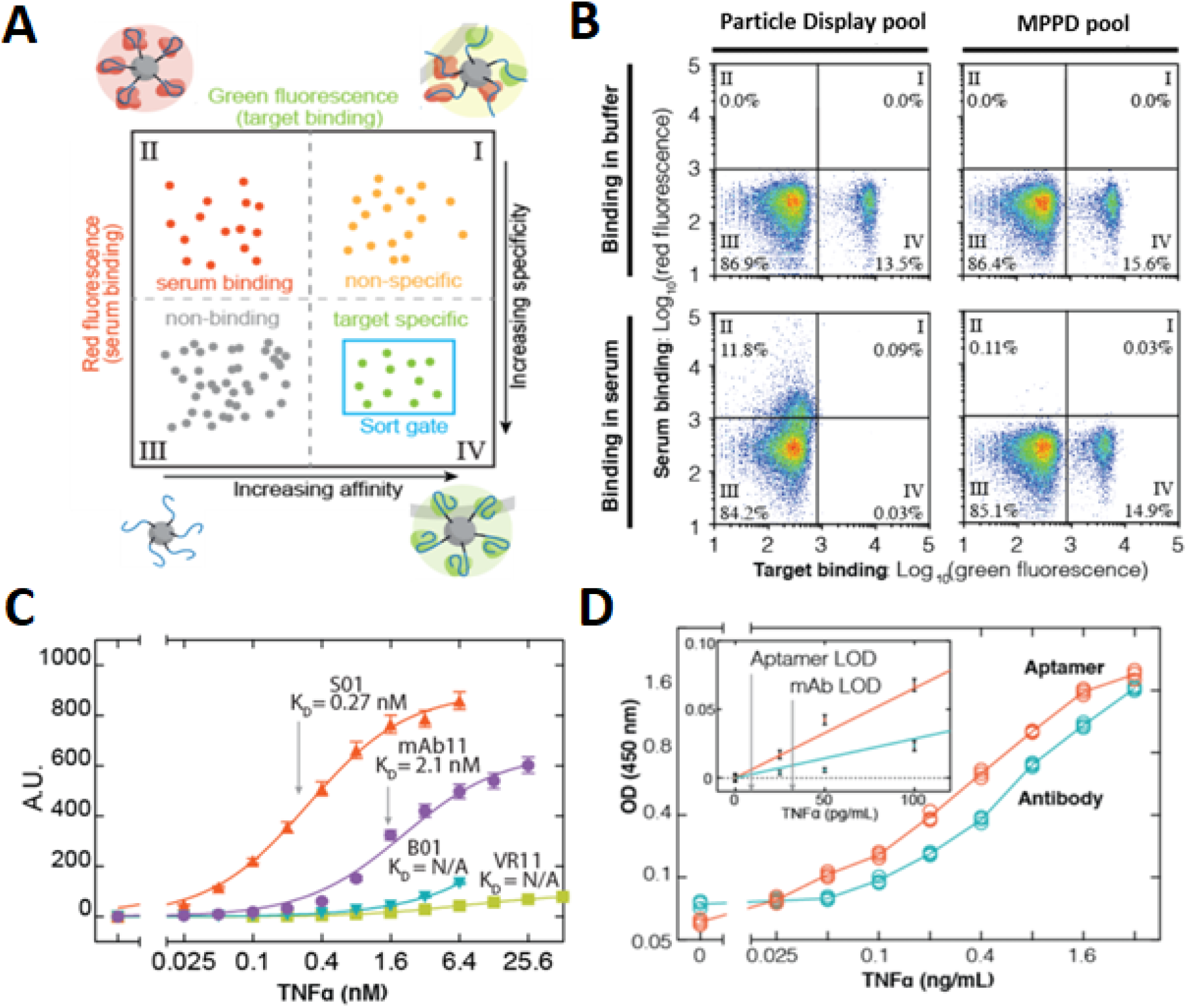
Multi-parameter particle display (MPPD) enables simultaneous screening of aptamer libraries for both affinity and specificity. **A**) In MPPD, the target is labeled with a green fluorescent tag, and non-target serum proteins are labeled with a red fluorescent tag. During FACS analysis, the goal is to isolate aptamer particles from quadrant IV, which exhibit both high target affinity (high green fluorescence) and high target specificity (low red fluorescence). **B**) Aptamers isolated via standard particle display in buffer (left) exhibit minimal binding to TNF-α in serum, indicating limited specificity, whereas those isolated via MPPD (right) perform equally well in both buffer and serum. **C**) TNF-α binding curves in serum for the top-performing MPPD (S01) and particle display (B01) aptamers relative to a previously published TNF-α aptamer (VR11) and a commercial TNF-α antibody (mAb11). **D**) S01 achieves a superior limit of detection to mAb11 in ELISA assays performed in serum.

When we analyzed the aptamer pools from these two screening experiments, it was clear that the dual-parameter selection yields far superior aptamers. The aptamers isolated via MPPD achieved equally strong and specific binding to TNF-α in both serum and buffer (**Figure 4B**, right panels), whereas aptamers generated via conventional particle display only exhibited strong binding to TNF-α in buffer (**Figure 4B**, top left), with no meaningful signal from TNF-α and extensive off-target binding in serum (quadrant II in **Figure 4B**, bottom left). We subsequently confirmed these findings by analyzing the highest-affinity aptamers from the buffer-only particle display (B01) and MPPD (S01) screens. Both S01 and B01 exhibited excellent affinity for TNF-α in buffer, considerably outperforming a previously reported aptamer (VR11) and a commercial antibody (mAb11) for the same target. However, B01 target affinity essentially disappeared in 10% diluted serum, presumably due to extensive off-target binding to interferent proteins (**Figure 4C**), while the affinity of S01 was virtually unchanged (*K*_*D*_ of 0.27 nM in serum versus 0.19 nM in buffer). Finally, we tested the performance of S01 in an ELISA, and found that our assay achieved a limit of detection (LOD) in serum that was more than three-fold lower than that of a commercial TNF-α ELISA kit (9.2 pg ml^−1^ versus 32 pg ml^−1^) (**Figure 4D**). These results demonstrate the clear advantages of using MPPD to actively select for aptamer specificity and affinity simultaneously, and highlight the risks associated with not incorporating specificity screening directly into the aptamer selection process.

### Expanding aptamer chemical diversity with click-PD

Although aptamers have successfully been generated against a wide range of targets, some targets remain stubbornly difficult, and there are data indicating that a sizeable proportion of the proteome might be inaccessible to natural DNA aptamers.^25,26^ Base-modified aptamers that incorporate non-natural chemical functional groups^27^ offer a greater diversity of chemical and structural properties than can be achieved with natural DNA or RNA, thereby expanding the spectrum of targets for which high-affinity and -specificity aptamers can be generated.^22,28–30^ For example, Gold and others developed a pioneering strategy for producing base-modified aptamers that tightly bind to a wide range of biomolecules that otherwise do not interact with natural DNA or RNA.^22,31^ The process of generating base-modified aptamers typically entails a laborious engineering process to generate polymerase variants that can incorporate and amplify chemically-modified nucleotides with minimal error rates.^32^ However, this may result in enzymes with impaired performance,^33,34^ and many chemical modifications are physically incompatible with polymerase-mediated incorporation.^33^ The Krauss^35,36^ and Mayer groups^37^ have overcome this problem with their ‘click-SELEX’ strategy, in which natural DNA nucleotides are substituted with an alkyne-modified alternative that can be readily coupled to an azide-modified functional group. However, this method is still subject to the above-mentioned limitations associated with conventional SELEX in terms of efficiency and the overall quality of the selected aptamer pool.

We have incorporated click-SELEX into our particle display technology, and this click-PD technique^13^ offers a general framework for the rapid and efficient selection of base-modified aptamers bearing virtually any chemical modification. Aptamer particles are generated as described above, but with alkyne-modified non-natural nucleobases incorporated into the PCR master mix during the emulsion PCR procedure (**Figure 5A**). These modified bases are readily incorporated by commercially-available polymerase enzymes, and can then be coupled to azide-tagged chemical functional groups via either copper-mediated or copper-free click conjugation. Some chemical groups need to be protected during click conjugation because they interfere with the reaction; during the single-strand generation step, these groups can be deprotected, after which the final base-modified DNA aptamer particle is ready for screening.

**Figure 5:**
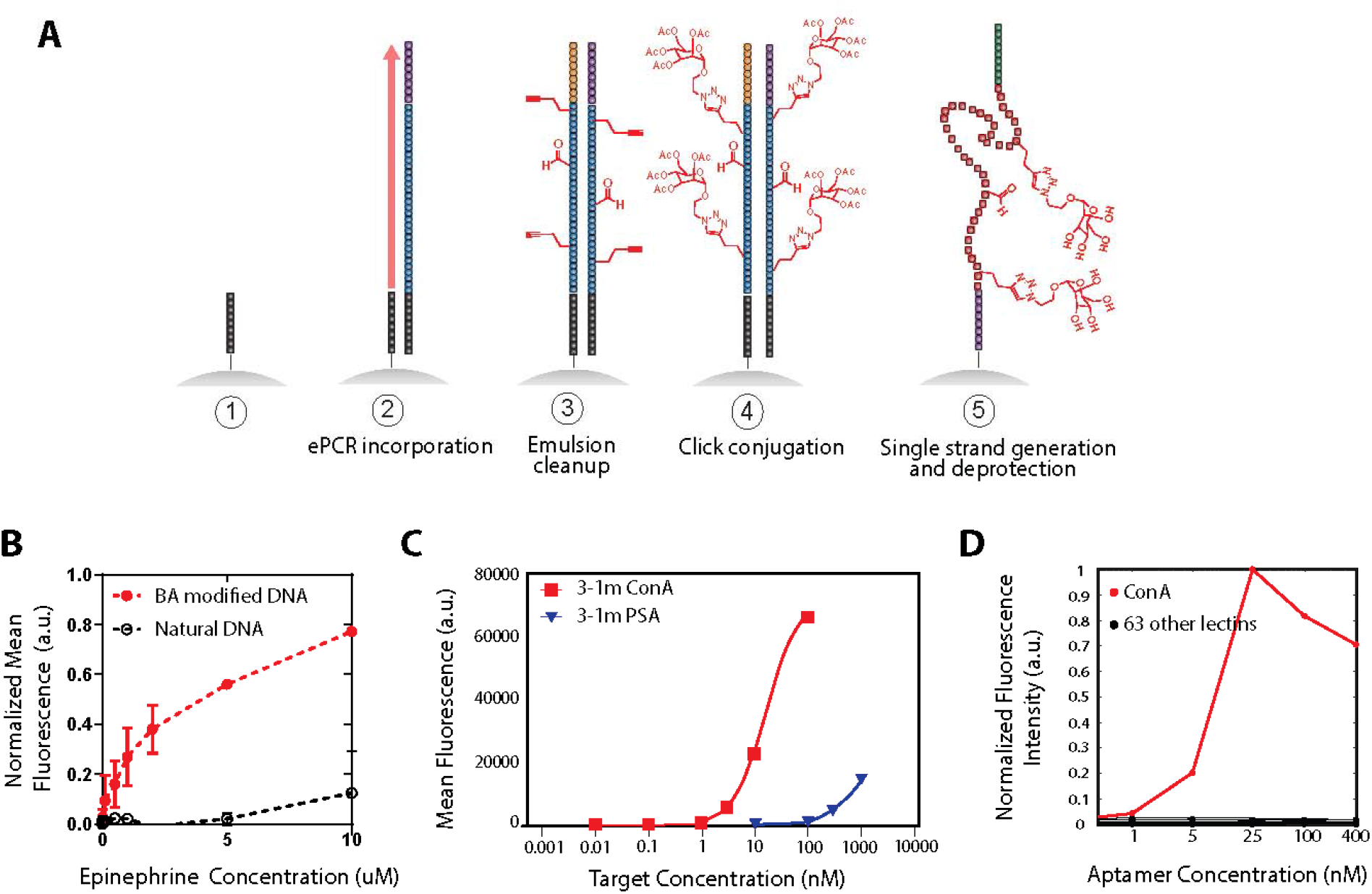
Click-PD enabled the generation of the first reported aptamer for epinephrine, as well as a highly specific Con A aptamer. **A**) Generation of base-modified aptamer particles in click-PD. 1) Forward primer-conjugated particles are generated via carbodiimide chemistry. 2) Monoclonal aptamer particles are generated via emulsion PCR with a mix of natural and non-natural dNTPs. The non-natural dNTPs include an alkyne handle that will be used during the click reaction. 3) The emulsion is broken and the particles – now decorated with double-stranded full-length aptamer sequences with alkyne groups – are collected. 4) Click conjugation is performed to attach azide-modified functional groups via either copper-catalyzed azide–alkyne cycloaddition (CuAAC) or strain-promoted azide-alkyne cycloaddition (SPAAC). 5) Single-stranded DNA is generated through a sodium hydroxide denaturation step. Protected chemical groups are deprotected during the same procedure. **B**) Flow cytometry-based binding assay for a boronic acid-modified epinephrine aptamer (red). When the modification is removed (black), binding is greatly reduced. **C**) Flow cytometry-based binding assay of a mannose-modified Con A aptamer. The aptamer is highly specific for Con A, and exhibits far weaker affinity for the non-target lectin PSA. **D**) An array-based binding assay testing the Con A aptamer against 63 other lectins shows that the aptamer is highly specific for Con A (red) and shows minimal binding to other known mannose binders.

As a demonstration, we performed a click-PD screen for epinephrine, a small-molecule neurotransmitter for which there were no published aptamers. We chose boronic acid as our modification, since this functional group forms a reversible covalent bond with a diol group^38–41^ found in epinephrine. Click-PD was initially developed to be compatible with a Huisgen copper-catalyzed azide-alkyne cycloaddition (CuAAC) click reaction. But since boronic acid interacts with copper, we instead performed a copper-free strain-promoted azide-alkyne cycloaddition (SPAAC) click reaction to attach the boronic acid modification to aptamers containing modified DBCO-dUTP. We performed one round of positive SELEX against epinephrine, and one round of negative SELEX to eliminate sequences that bind to the particles or the fluorescein isothiocyanate (FITC) dye used to label the epinephrine. We then performed four rounds of click-PD with FITC-labeled epinephrine, followed by HTS to identify aptamer sequences enriched over the course of screening. A flow cytometry bead-based assay revealed that our top boronic acid-modified aptamer had a *K*_*D*,*eff*_ of 1.1 μM; when the boronic acid modification was removed, the aptamer showed negligible binding to epinephrine (**Figure 5B**). By way of comparison, aptamers for small molecules generally exhibit modest affinity, with *K*_*D*_ in the 10– 100 μM range,^42^ demonstrating the robust performance of our boronic acid-modified aptamer.

As with MPPD, click-PD can be used to simultaneously select for base-modified aptamers that achieve both high affinity and specificity for their target. To demonstrate this capability, we screened for glycomimetic aptamers that can bind strongly and selectively to a particular lectin, concanavalin A (Con A). Lectins are a large class of carbohydrate-binding proteins that often exhibit high structural homology, and thus pose a considerable challenge in terms of generating affinity reagents that achieve high specificity^43^. We generated a large library of mannose-modified aptamers in order to identify ligands for Con A, which is widely used to study the molecular mechanisms of protein-carbohydrate interactions.^44,45^ We generated particles that display aptamers incorporating a modified dUTP with an alkyne handle, and then performed a CuAAC reaction to conjugate mannose groups to the DNA. We used a two-color approach to simultaneously select for Con A affinity and specificity; as a non-target competitor molecule, we selected *Pisum sativum* agglutinin (PSA), a lectin with considerable structural homology to Con A.46 After three rounds of FACS to isolate aptamers with high binding to Con A and low binding to PSA, we identified 3-1m, a base-modified aptamer that binds Con A with a *K*_*D*_ of 17 nM and remarkable target specificity. Not only did 3-1m have orders of magnitude higher affinity for Con A than PSA (**Figure 5C**), but it also showed no or minimal binding (< 30% of maximum signal) to an array displaying 63 other lectins (**Figure 5D**). This strong affinity and specificity also conferred potent biological activity, and we demonstrated that 3-1m could inhibit the hemagglutination reaction with three-fold greater potency than the best Con A inhibitor described to date.^47^ Thus, click-PD makes it possible to screen for base-modified DNA aptamers featuring diverse chemical groups that simultaneously achieve high affinity and specificity.

### High-throughput characterization of base-modified aptamers with N2A2

The characterization of the aptamers that result from selection has remained a highly resource-intensive step in the aptamer discovery process. This is because aptamers are typically characterized in terms of their binding affinity and specificity either individually in a serial fashion or at very low throughput in parallel. Commonly-used assays for measuring *K*_*D*_, such as flow cytometry and surface plasmon resonance, offer a maximum throughput of just 10–100 sequences per experiment, compared to the 10^6^–10^7^ candidates one would ideally want to test after HTS analysis of an aptamer pool. And since these screening methods are also time-consuming to perform, rare optimal aptamer candidates are likely to be missed. We have previously described the quantitative parallel aptamer selection system (QPASS) as a solution to this problem.^48^ QPASS couples microfluidic selection and HTS analysis with microarrays to measure the binding affinity of thousands of aptamer candidates simultaneously, and has demonstrated the capacity to rapidly identify aptamer candidates that perform optimally in a variety of biological samples and assay conditions. Several groups have also demonstrated efficient screening of natural DNA and RNA libraries by repurposing HTS instruments to achieve parallel characterization of protein binding for millions of sequences in parallel within an Illumina flow-cell.^49–52^ However, none of these techniques is suitable for the functional characterization of base-modified aptamers, which remains a low-throughput process.

We have recently developed a more sophisticated platform for high-throughput aptamer analysis known as N2A2. Our goal was to design a system that can perform synthesis, screening, and binding characterization of chemically base-modified aptamers in a single integrated workflow.^53^ N2A2 employs a similar concept to the HTS-based systems described above for characterizing natural RNA or DNA sequences, wherein minor modifications to a relatively low-cost benchtop sequencing instrument – the Illumina MiSeq – enable us to characterize binding of ~10^7^ base-modified aptamers in parallel in a semi-automated fashion. As with click-PD, N2A2 employs a click chemistry-based modification approach that enables efficient incorporation of virtually any chemical functional group with commercially available polymerase enzymes. This makes it straightforward to screen for the chemical modification that results in the highest affinity and specificity aptamer for a given target—a process that otherwise relies heavily on educated guesses.^54^ N2A2 is also suitable for the characterization of natural DNA aptamer pools, and can achieve massively parallel characterization of aptamer binding at a throughput that is several orders of magnitude higher than QPASS (~10^7^ versus ~10^4^ sequences).

The N2A2 workflow (**Figure 6Error! Reference source not found.**) begins with sequencing of a DNA library. During the paired-end turnaround and second read of the sequencing process,^55,56^ the DNA clusters are converted into base-modified aptamer clusters. As with click-PD, we make use of commercial polymerases that can efficiently incorporate modified bases with click handles. The flow-cell is then incubated with fluorescently-labeled protein at various concentrations to screen for affinity and specificity, with intensity information collected for each cluster. These sequencing and screening data are processed to generate a phenotype-genotype linked map of all the clusters.

N2A2 offers a versatile platform for tackling aptamer development problems that were challenging to solve with existing technologies. The ease with which different modifications can be interchangeably incorporated into a given aptamer library makes it straightforward to test how different chemical functional groups affect target binding. As a demonstration of this, we performed two N2A2 screens for the vascular endothelial growth factor (VEGF) protein with the same pool of candidate aptamers modified either with tyrosine or tryptophan. We determined that tryptophan-modified aptamers consistently showed higher binding for VEGF, and identified a tryptophan-modified VEGF aptamer with a *K*_*Deff*_ of 2.8 nM, which is 6.6-fold higher affinity than a previously published natural DNA aptamer^57^ (**Figure 6A**).

**Figure 6.**
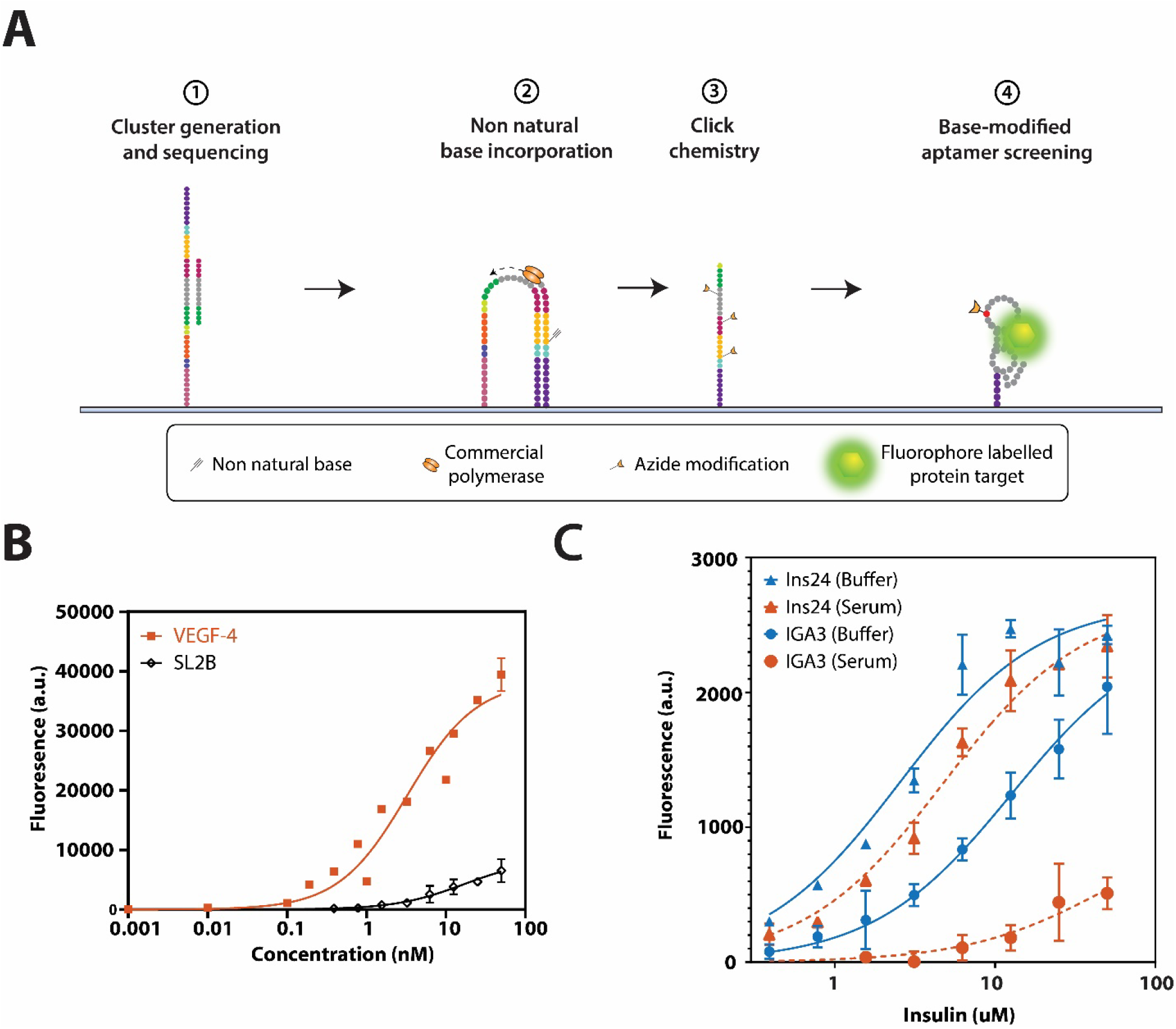
A) Overview of N2A2. 1) The workflow begins with sequencing of a DNA library within the MiSeq flow-cell. 2) Next, the DNA clusters are converted into base-modified aptamer clusters via 3) click chemistry. 4) Finally, the aptamers are incubated with fluorescently-labeled protein to screen for affinity and specificity, with intensity information collected for each cluster. B) Flow cytometry characterization demonstrates that our tryptophan-modified aptamer VEGF-4 (orange) shows greater affinity for VEGF than a previously described aptamer, SL2B (blue). C) A flow cytometry-based binding assay for the previously published insulin aptamer IGA3 (circles) and the N2A2-identified aptamer ins24 (triangles) in buffer (blue, solid lines) and 1% human serum (orange, dotted lines). Only ins24 retains insulin binding in serum.

N2A2 is also capable of performing direct screening in complex sample mixtures, enabling the generation of base-modified aptamers with excellent specificity for their target. To demonstrate this, we screened a library of phenylalanine-modified aptamers against the peptide hormone insulin in buffer and 1% diluted human serum. We identified a number of aptamers that exhibit excellent target affinity and specificity, and further characterized several of these sequences in a flow cytometry assay. The top aptamer from this screen, Ins24, exhibited consistent target affinity in both buffer and serum, demonstrating the effectiveness of N2A2 in selecting for specificity. In contrast, a previously published natural DNA aptamer, IGA3,^58^ exhibited lower affinity than Ins24 in buffer and entirely lost its ability to bind insulin in the complex background of diluted serum (**Figure 6B**).

## Conclusion

In this Account, we have chronicled our lab’s progress in overcoming many challenges impeding the isolation of high-quality aptamers via SELEX. First, we developed particle display to dramatically improve the efficiency with which high-affinity aptamers can be enriched in a single round of screening. Next, we extended the particle display technology with MPPD, which allows us to simultaneously screen for aptamers that exhibit both excellent affinity and specificity for their target. We also adapted particle display for use with non-natural bases with our click-PD technology, which allows researchers to expand the range of targets accessible to aptamers by introducing greater chemical diversity into their libraries. Finally, we have developed a high-throughput solution for characterizing aptamer affinity and specificity with the N2A2 platform, which makes it possible to generate, screen, and characterize the binding properties of tens of millions of base-modified aptamers in a single experiment.

These technological advances greatly extend the utility of aptamers, but there are still additional challenges to overcome. Foremost among these is finding ways to access the sequence space that is currently physically unexplorable: a library with a 40-nt random region has 10^24^ possible sequences, but realistically we can only screen 10^15^ in an experimental setting. Protein engineers have utilized machine learning (ML) techniques to computationally address this problem in the context of enzyme^59,60^ and antibody^61,62^ engineering, and the aptamer field has also begun to develop equivalent ML solutions. One early example is closed-loop aptameric directed evolution (CLADE)^63^, an ML tool that utilizes data from microarray-based fitness assays to perform additional selection and mutation of aptamer sequences *in silico*. Bashir *et al.* have continued this work by developing the ML-guided Particle Display methodology (MLPD)^64^, in which they were able to predict high affinity aptamers for the neutrophil gelatinase-associated lipocalin (NGAL) protein based on the sequencing results from a particle display screen, as well as the minimal motifs that determine aptamer binding. Further integration of ML into the aptamer development toolbox will require the generation of sufficient quantities of high-quality training data—including both positive and negative examples—as well as a determination of how much training data is necessary to routinely generate robust predictions. To date, HTS data from previous aptamer selections have been used as training datasets, but these sequence data are not directly coupled with data pertaining to functional performance, and this lack of robust experimental data for training and classifying aptamer-target interactions has delayed development of aptamer ML methods. We believe N2A2 offers a powerful solution here, with the capacity to generate comprehensive mutational maps of aptamer sequences that can be directly linked to their effects on functional performance.

## Acknowledgements

The authors would like to thank Alex Rangel and Brian Young for their constructive and valuable input and comments in preparing this Account. This work was supported by the National Institutes of Health (R01 GM129312, OT2OD025342), Chan-Zuckerberg Biohub, and the Bill and Melinda Gates foundation.

